# Tools for engineering coordinated system behaviour in synthetic microbial consortia

**DOI:** 10.1101/231597

**Authors:** Nicolas Kylilis, Guy-Bart Stan, Karen M Polizzi

**Affiliations:** Department of Bioengineering, Imperial College London, London SW7 2AZ, United Kingdom; Department of Life Sciences, Imperial College London, London SW7 2AZ, United Kingdom; Centre for Synthetic Biology and Innovation, Imperial College London, London SW7 2AZ, United Kingdom

**Keywords:** Synthetic biology, Quorum sensing, Synthetic microbial consortia, Cell-to-cell communication, Characterisation, Computer aided design

## Abstract

Advancing synthetic biology to the multicellular level requires the development of multiple orthogonal cell-to-cell communication channels to propagate information with minimal signal interference. The development of quorum sensing devices, the cornerstone technology for building microbial communities with coordinated system behaviour, has largely focused on reducing signal leakage between systems of cognate AHL/transcription factor pairs. However, the use of non-cognate signals as a design feature has received limited attention so far. Here, we demonstrate the largest library of AHL-receiver devices constructed to date with all cognate and non-cognate chemical signal interactions quantified and we develop a software tool that allows automated selection of orthogonal chemical channels. We use this approach to identify up to four orthogonal channels *in silico* and experimentally demonstrate the simultaneous use of three channels in co-culture. The development of multiple non-interfering cell-to-cell communication channels will facilitate the design of synthetic microbial consortia for novel applications including distributed bio-computation, increased bioprocess efficiency, cell specialisation, and spatial organisation.

---

Synthetic biology research and applications to date have mostly been focused on the engineering of homogeneous or monoclonal designer cell populations to perform functions ranging from biocomputation^1^, to bioproduction of biomaterials^2^ and chemicals^3^, to biosensing^4,5^ among others. *De novo* implementation of genetic circuits of increasing complexity and size in living cells is an important challenge in synthetic biology. Obstacles include unwanted interactions between genetic parts^6^, restrictions on the number of available genetic parts^1^, limits to the size of DNA that can be transformed into the host cell and metabolic burden to the host chassis^7^. The use of consortia of organisms could potentially alleviate these bottlenecks if the functionality of the genetic circuit can be distributed among different cell populations^8^. Additionally, the use of synthetic consortia could expand the capabilities and applications of synthetic biology by enabling compartmentalisation^9^, cell specialisation^10^, parallel bio-computation^11^, increased bioprocess efficiency^12^ and spatial organisation^13^.

A vital component for the engineering of synthetic consortia are devices that enable communication between the different populations so as to coordinate their behaviour. Quorum sensing systems, especially those based on small molecule acylserine homolactones (AHL), have become the favoured technology for engineering cell-to-cell communication^14^ because of their simple genetic architecture. AHL is produced enzymatically by the expression of a single enzyme, e.g. the Acyl-homoserine-lactone synthase protein that is the product of the *luxI* gene^15^. AHL molecules can freely diffuse in the intracellular and extracellular environment. Intracellularly, AHLs bind transcription factor proteins, which results in an activated complex that can bind a quorum sensing promoter to initiate transcription of downstream genes^16^. The most frequently used quorum sensing parts, the LuxR protein and its cognate Plux promoter have been assembled into a AHL-receiver device that has been extensively characterised in terms of its input-output response, dynamic performance upon induction, specificity to other AHL molecules, and evolutionary reliability^17^. Various studies have sought to increase the number of available AHL communication modules with functional devices built from the biological components of the las^18^, tra^18^, rpa^18^, rhl^19^, cin^19^, and esa^20^ quorum sensing systems. Nevertheless, quorum sensing devices can exhibit various degrees of crosstalk either in the form of AHL molecules binding to non-cognate transcription factors (chemical crosstalk) or transcription factors binding to non-cognate promoters (genetic cross-talk)^13^. The degree of orthogonality between designed AHL communication modules can be quantified by high-throughput screening as demonstrated for a set of designed AHL-receiver devices of the lux, las, rpa, and tra quorum systems^18^. For this set of devices only the tra and rpa devices were determined to be completely orthogonal for both chemical and genetic crosstalk^18^. However, a number of strategies have been demonstrated for minimising crosstalk, which include modulating the expression levels of the transcription factor that influence the response function of the device, and quorum sensing promoter engineering^13^.

In this research, we aimed to increase the number of available tools for the design of microbial communities for novel applications. Initially, we designed and constructed AHL-receiver devices from components of the rhl, lux, tra, las, cin and rpa quorum sensing systems and characterised their input/output behaviour at the individual cell level. Next, we characterised the functionality of these AHL-receiver devices in the presence of non-cognate AHL inducers. In doing so, we created the largest characterized library of quorum sensing devices with mapped chemical crosstalk interactions. This large database and the associated algorithm that we developed to automate the selection of compatible and orthogonal AHL communication channels facilitate the design of synthetic microbial consortia. We demonstrate the power of this approach by experimentally validating one of the algorithmically proposed *in silico* designs for the specific control of gene expression through three non-interfering AHL communication channels in a polyclonal *E. coli* co-culture.

## RESULTS

### Design and characterisation of AHL-receiver devices

To expand the pool of well-characterised cell-to-cell communication tools for engineering microbial consortia, synthetic AHL-receiver devices were constructed that employed components from six quorum sensing systems (Figure 1). The engineered systems incorporated genetic part elements from the lux system^21^ (*Vibrio fisheri*), the rhl^22,23^ and las^24,25^ systems (*Pseudomonas aeruginosa*), the cin^26^ system (*Rhizobium leguminosarum*), the tra^27^ system (*Agrobacterium tumefaciens*) and the rpa^28^ system (*Rhodopseudomonas palustris*). The cognate AHL-inducer signal molecule for each system is indicated in Figure 1. The AHL-receiver devices share an identical genetic architecture and were cloned into the pSB1C3 vector backbone for propagation in cells. For *in vivo* characterisation of the AHL-receiver devices, a GFP reporter was cloned downstream of the quorum sensing promoters and cells were transformed with the resulting composite device plasmids. The transformed cells were cultured, induced using the cognate AHL, and the resulting cell fluorescence was assayed by flow cytometry. All the AHL-receiver devices functioned as designed as demonstrated by the increase in cell fluorescence at higher AHL concentration (Figure 2A) with homogeneous GFP expression output at all AHL-inducer concentrations. This is despite differences in the tail length of inducer molecules and the presence of additional functional groups that may affect the permeability of the cell membrane to the inducer^29^. Many applications in synthetic biology require careful modulation of intracellular gene expression levels in response to inducer concentrations. However, a number of inducible systems frequently used in genetic circuit design exhibit an on or off response at the individual cell level despite the appearance of a graded response at the population level^30^. The six devices analysed here avoid this and can therefore be used to tune genetic expression levels over a wide range.

**Figure 1:**
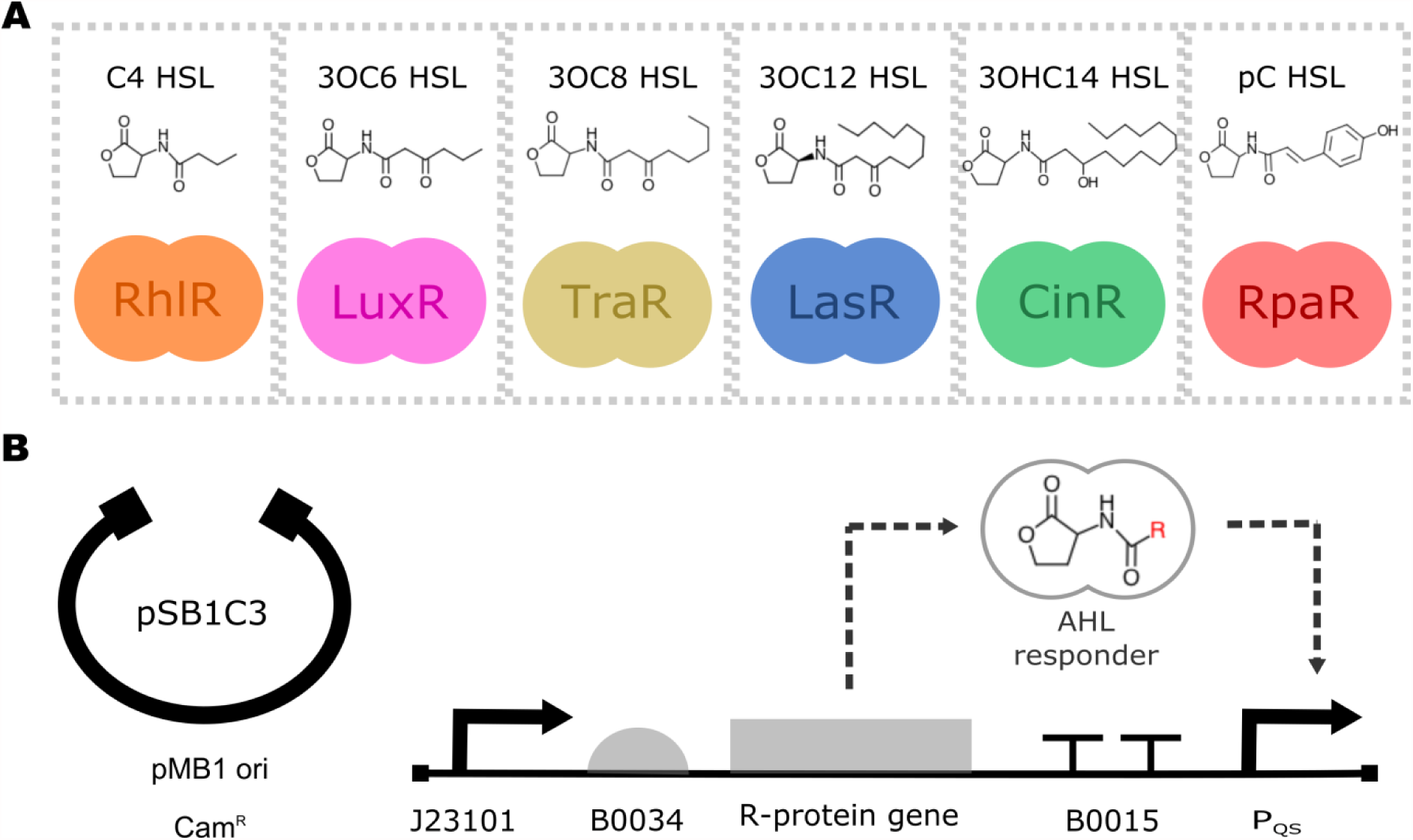
Library of AHL-receiver devices. **(A)** Quorum sensing systems with schematics of the transcription factor regulator protein and chemical structures of the cognate AHL ligand. (**B**) Vector backbone and genetic architecture of AHL-receiver devices.

**Figure 2:**
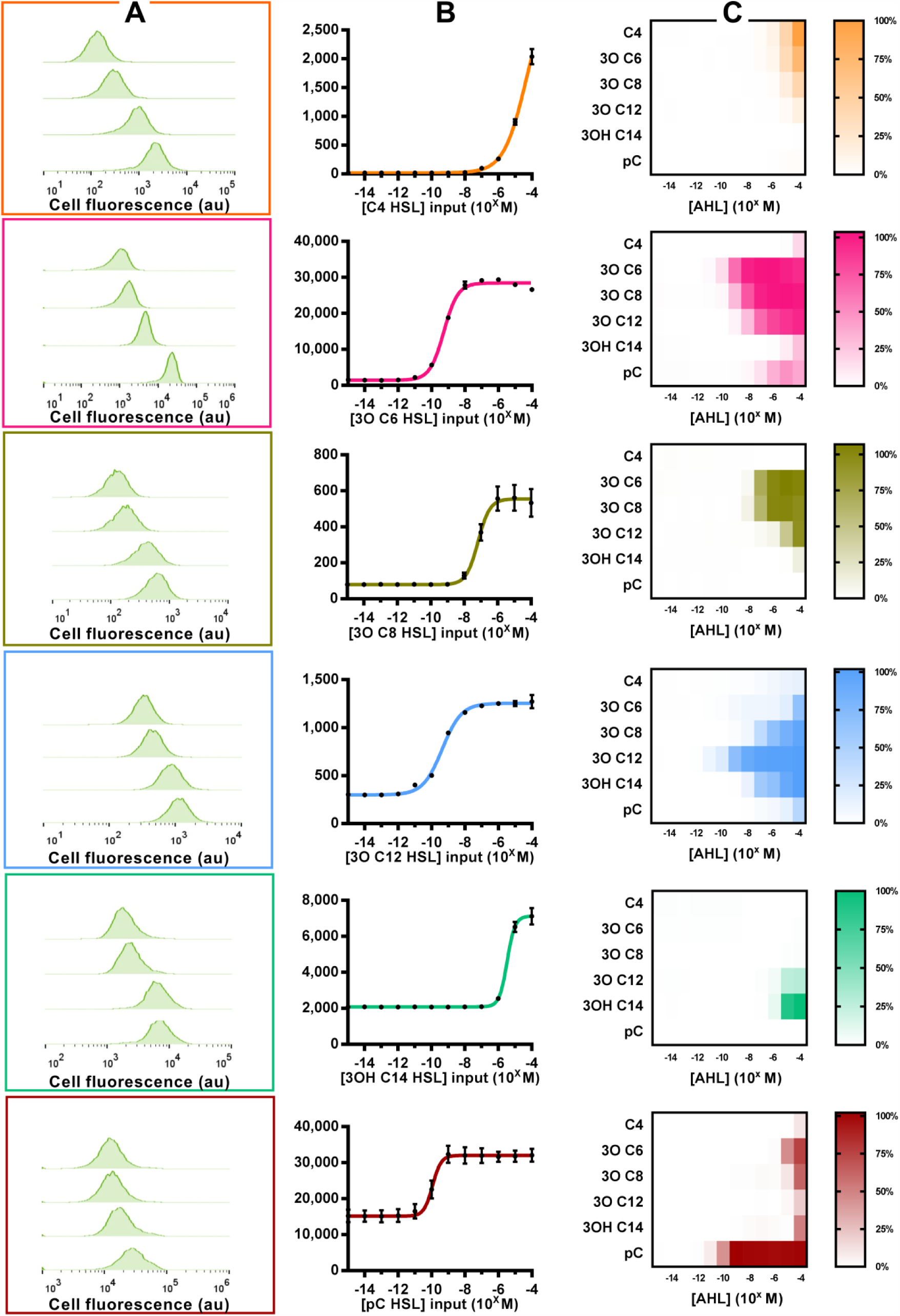
Characterisation of AHL-receiver devices. **(A)** Cell fluorescence from flow cytometry of AHL-receiver composite devices/AHL inducers cognate pairs at selected inducer concentrations. (**B**) Input/output functions of AHL-receiver devices, i.e. GFP output (au) against cognate AHL inducer concentrations derived when mean fluorescence data were fitted with a 4-parameter logistical function model (Equation 1). Error bars represent the standard error of the mean of 3 biological repeats. (**C**) Heat maps of normalised GFP output from AHL-receiver composite devices induced with cognate and non-cognate AHL inducer concentrations.

The input/output function of the AHL-receiver devices was determined by fitting the data (Figure 2B) with a 4-parameter logistical curve model (Equation 1). This model takes the form of:

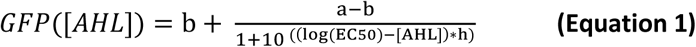

where a is the maximal GFP output, b is the basal GFP output, [AHL] is the AHL inducer concentration, GFP is the response of the device for that inducer concentration, EC50 is the inducer concentration that results in half maximal activation of the device, and h is proportional to the value of the steepest slope along the curve (Hill coefficient) that indicates the responsiveness of the device to the input. To facilitate the computer-aided-design of genetic circuits for engineering microbial consortia systems when using these devices, the parameters values for the transfer function of each device are provided in the Supplementary Information.

### Characterisation of chemical signal crosstalk for non-cognate AHL-receiver devices

Next, we sought to investigate the chemical crosstalk of the AHL-receiver devices as this can severely limit the engineering of synthetic consortia requiring the use of multiple chemical communication channels. The characterisation of the GFP output for all six AHL-receiver devices was carried out as previously for each of the 5 non-cognate AHL inducers, thus creating the largest library of AHL-receiver devices with all their chemical signal interactions quantified to date. The experimental results are presented in Figure 2C in the form of heat maps of normalised GFP output for each device.

These results show that each device exhibited a distinct profile of crosstalk interaction and all devices were effectively activated by non-cognate AHL molecules. Amongst the devices, the rpa system exhibited the least propensity for activation by non-cognate AHL molecules in comparison to its cognate AHL inducer. This likely stems from the distinct molecular structure of the p-coumaryl HSL inducer compared to the rest of the AHL molecules. Despite the apparent chemical crosstalk when considering operation over the whole range of AHL induction, it was clear that the use of particular combinations of AHL-receiver devices operated within restricted AHL inducer concentration regimes could allow the simultaneous use of multiple orthogonal communication channels.

### Characterisation of the relative activity of devices to simplify coupling to other technologies

To facilitate the re-use of the AHL-receiver devices demonstrated here in combination with other synthetic biology devices and systems, the basal and maximal expression levels were calibrated against a reference promoter (Figure 3B). Similar to previous work^31^, the *in vivo* activity of the J23101 promoter was set as the reference standard with relative promoter activity of 1, and expression characteristics of the engineered AHL-receiver devices were calibrated according to this reference. Additional Andersen promoters were included to facilitate benchmarking against lower strength promoters where necessary. The engineered systems were determined to vary widely both in relative basal and maximal expression strength (Figure 3C). This is an important consideration for their integration with downstream modules in engineered systems. Furthermore, their documented, widely varied relative expression strength provides a broad design space for genetic circuit design.

**Figure 3:**
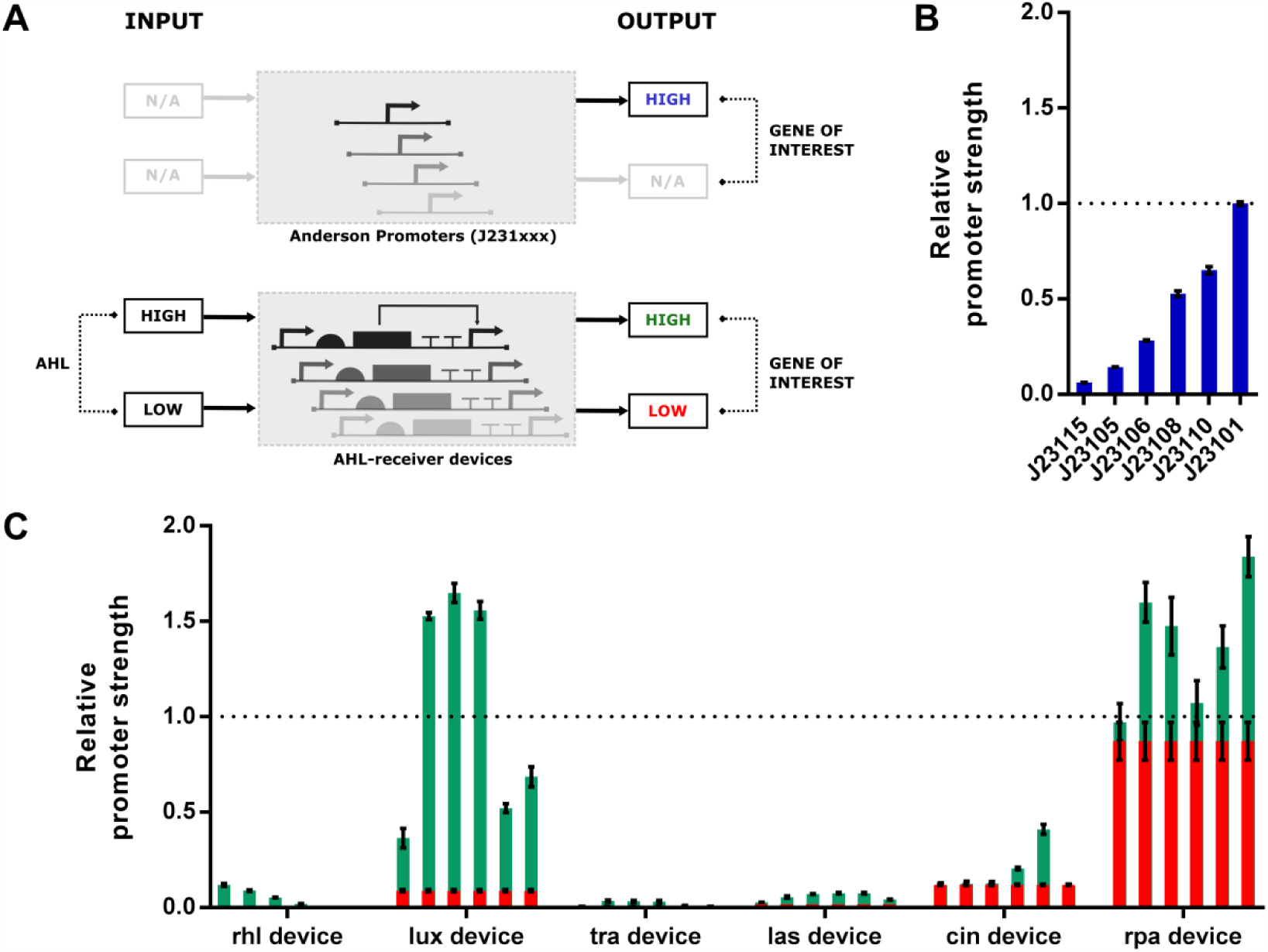
Relative promoter strengths of AHL-receiver devices. **(A)** Schematic of possible states of characterised genetic devices. Constitutive promoters of the Anderson promoter library exhibit just one output state of high signal in the absence of any input signal. Conversely, the AHL-receiver devices can operate in two states: at low input they exhibit a low output; while at high input they exhibit high output. (**B**) Bar chart of relative promoter strength of constitutive Anderson promoters and (**C**) inducible AHL-receiver devices. The dotted line indicates promoter strength of the reference standard promoter Bba_J23101. The chart is color-coded according to the states of the devices as indicated in the schematic in (A). Error bars indicate the standard error of the mean of three biological repeats.

### Software tool for automated identification of orthogonal chemical communication channels

To facilitate the genetic circuit design and manipulation of microbial consortia, a computational algorithm was developed that automatically identifies combinations of AHL-receiver devices and AHL inducers that allow for “effective orthogonality” according to user-defined specifications. The program uses the fitted model parameters of the AHL-receiver devices with cognate and non-cognate AHLs to identify suitable combinations of AHL-receiver devices and AHL inducer concentrations that behave orthogonally within a given experimental design. The user can specify activation thresholds for specific gene expression, crosstalk thresholds and the number of chemical communication channels required.

To demonstrate the utility of this tool, the algorithm was used to identify suitable systems that would allow for a number of chemical communication channels with minimal signal crosstalk in a hypothetical engineered consortium. Initial specifications focused the identification of two quorum sensing systems that would allow for more than two-fold activation of specific gene expression, while exhibiting less than two-fold non-specific activation of gene expression (crosstalk). The program was able to identify several combinations of AHL-receiver pairs that met the specifications. Amongst the identified systems were pairs of systems that utilised their cognate AHL inducers (Figure 4A, top), pairs of systems that utilised non-cognate AHL-inducers (Figure 4A, middle) and pairs of systems that utilised a combination of cognate and non-cognate AHL inducers (Figure 4A, bottom). In a further example, stricter input specifications were set for a specific activation threshold of more than ten-fold and a crosstalk threshold of less than three-fold for two communication channels. The algorithm identified a system consisting of the rhl device with C4 HSL [1×10^−6^ – 1×10^−5^ M] and the lux device with 3O C6 HSL [1×10^−9^ – 1×10^−7^ M] (Figure 4B) that meets the pre-defined orthogonality criteria. Furthermore, the algorithm was able to identify several systems for the simultaneous use of three orthogonal chemical communication channels (an example presented in Figure 4C), and even a system of four channels with minimal signal crosstalk (Figure 4C) with more than two-fold specific activation and less than two-fold crosstalk.

**Figure 4:**
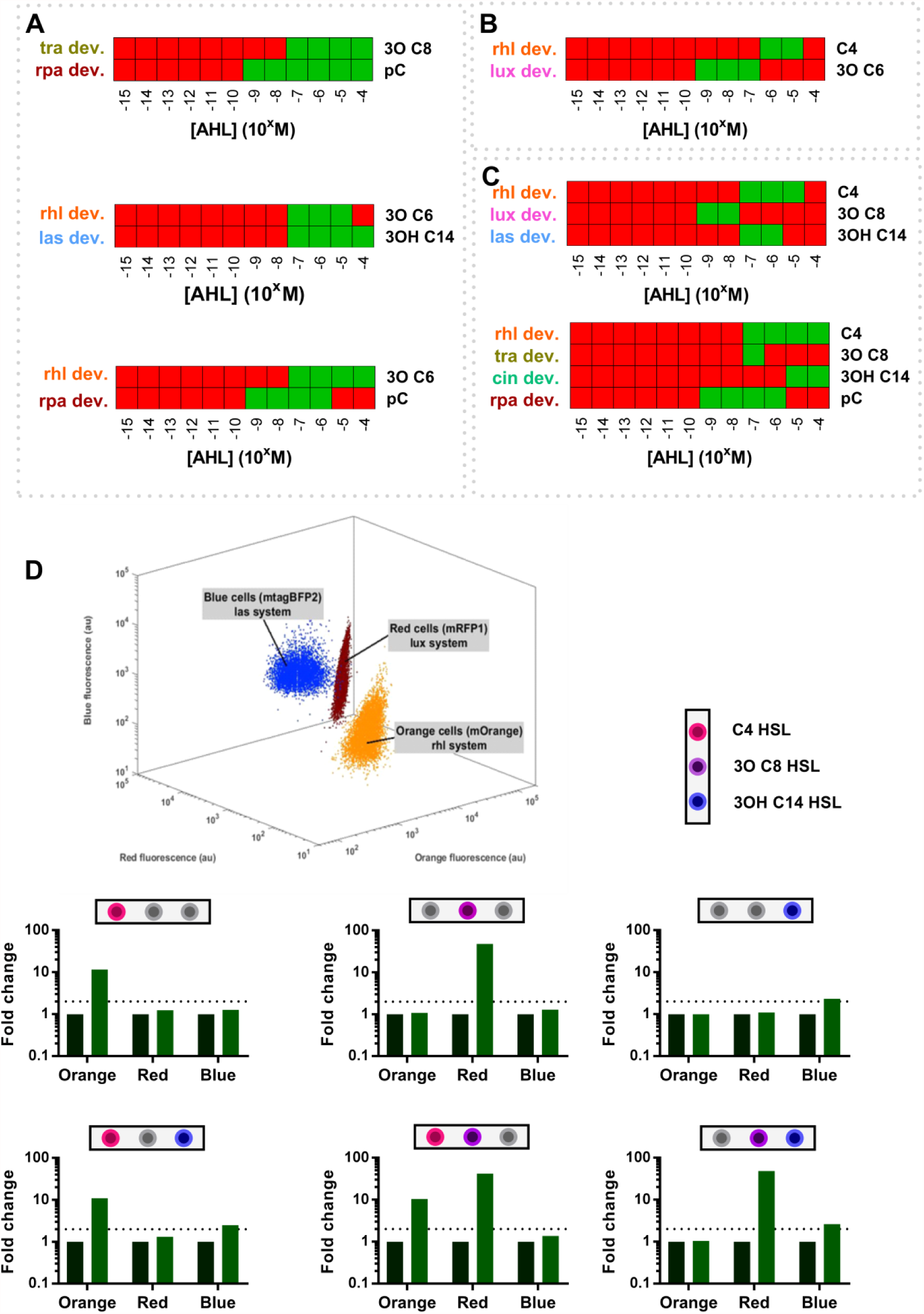
Computer-aided-design of synthetic microbial consortia with orthogonal communication channels. **(A-C)** Identified combinations for orthogonal AHL-chemical communication channels as provided by our computer-aided-design software. For each panel, red indicates an AHL-inducer concentration which does not meet at least one of the user-defined specifications, while a green panel indicates an AHL-inducer concentration which meets all the specifications. (**A**) Systems of two communication channels designed with the following specifications: two-fold gene activation and less than two-fold crosstalk. (**B**) System of two communication channels designed with the following specifications: more than ten-fold gene activation and less than three-fold crosstalk. (**C**) System of three communication channels (top) and four communication channels (bottom) designed with the following specifications: more than two-fold gene activation and less than two-fold crosstalk. (**D**) Co-culture of three *E. coli* populations predicted to be able to control gene expression using three orthogonal chemical communication channels (panel C top); differentiated by flow cytometry (3D-pot) for mOrange (orange cell population), mRFP1 (red cell population) or mtagBFP2 (blue cell population) fluorescent protein production. The calculated fold-change in GFP output for each AHL-chemical channel is indicated below (D). The dotted line in the bar charts represents the user-specified threshold value for gene activation and crosstalk signal.

To experimentally validate the predictive capabilities of the software tool, we implemented the design for the system with three orthogonal communication channels presented in Figure 4C (top panel). The suggested system was implemented in genetically modified *E. coli* strains using the rhl, lux and las composite devices with a GFP reporter module. The engineered strains were also transformed with plasmid constructs encoding constitutive expression of mOrange, mRFP1 or mtagBFP2 fluorescent protein devices to enable bacterial strain differentiation when in co-culture (Figure 4D 3D-plot). The co-culture system was tested for the presence of orthogonal communication channels by the addition C4 HSL [1×10^−5^M], 3O C8 HSL [1×10^−8^M] and 3OH C14 HSL [1×10^−7^M] in all single and dual input combinations with the change in cell fluorescence being recorded by flow cytometry. The experiment results showed that the calculated fold change in GFP output was more than two-fold for specific gene expression and less than the two-fold threshold set for crosstalk gene activation in all three chemical communication channels (Figure 4D), in complete agreement with the prediction of the computer-aided-design algorithm.

## CONCLUSIONS

In this work, we describe the construction of the largest library of AHL-receiver devices reported to date. This library consists of six AHL-receiver devices constructed using genetic parts mined from various microbial species. The input/output functions of the devices were measured for their cognate and non-cognate AHL-inducer molecules, and characterised in terms of their basal expression, maximal expression, fold activation, input signal EC50 concentration for half-activation and device sensitivity. Such metrics are important in assessing the device suitability for use in genetic circuit designs. For example, the fold activation metric can be of significance when incorporating these devices with downstream processes that require overcoming specific thresholds for activation or inhibition, such as in logic gate designs^1,32^. At the same time, the EC50 value can be of importance for engineering bacterial populations that respond to cell population densities^33^.

Additionally, both cognate and non-cognate pairs of AHL-inducer/AHL-receiver devices have been shown to be able to activate gene expression at different levels of expressions. This behaviour, termed as chemical crosstalk, can complicate the design of synthetic microbial consortia. Here, through the characterisation of a total of 36 synthetic quorum systems (cognate and non-cognate pairs) and the development and use of a computer-aided-design tool, we were able to identify ranges of AHL-inducer concentrations that delineate orthogonal chemical communication. This enables the simultaneous use of our AHL-receiver devices for the engineering of microbial consortia. We used our approach to identify a potential system using up to four orthogonal channels simultaneously. We experimentally validated this approach by engineering a polyclonal co-culture capable of controlling gene expression using three non-interfering AHL communication channels. Additionally, the computer-aided-design (CAD) tool (see Supplementary Information for the code) is designed to aid the identification of a user-specified number of orthogonal communication channels from a library of characterised AHL-receiver devices. This CAD software is easily adaptable to accommodate specific user-defined constraints and number of AHL-devices, which can be particularly useful in enabling consortia designs of increasing complexity as the number of characterised quorum systems expands.

To fully realise the potential of engineered microbial consortia, further research could provide methodologies and sets of design rules to further expand the chemical design space for orthogonal communication and control of gene expression in multicellular systems. Beyond effective cell-to cell communication technologies, another important aspect is designs that allow robust maintenance of cell populations in co-cultures in order to ensure the long-term co-existence of different engineered species. Development of effective design strategies that enable the rational engineering of synthetic microbial consortia will lead to novel application areas^34^, and further tools for understanding natural ecosystems^35,36^. Additionally, the engineering of microbial consortia would provide a foundational framework for tissue/organ engineering^37,38^, and transformative medical therapeutic applications.

## METHODS

### Strains, plasmid constructs and chemicals

Experiments were carried out with *E. coli* cells Top10 strain transformed with plasmid constructs. Plasmid constructs were assembled to comply with the BioBrick RCF [10] standard using a variety of molecular cloning techniques. Biological parts were acquired from the registry of Standard Biological parts or from chemical synthesis. DNA sequences of assembled AHL-receiver devices are available in the Supplementary Information. AHL inducers utilised were purchased as follows: N-Butyryl-DL-homoserine lactone (09945 Sigma), 3-oxohexanoyl-L-homoserine lactone (K3007 Aldrich); N-(3-Oxododecanoyl)-L-homoserine lactone (O9139 Sigma) and N-(3-Hydroxytetradecanoyl)-DL-homoserine lactone (51481 Sigma). AHLs stock solutions were made using 100% DMSO as a solvent.

### Cell cultures and AHL induction

*E. coli* Top10 cells transformed with plasmid constructs were cultured for *in vivo* GFP expression measurements as follows: Overnight culture of transformed cells were diluted 1:100 into LB medium supplemented with 34 μg/mL chloramphenicol and grown for 3 hours at 37°C with shaking at 250rpm. Cultures were diluted to an OD_600_ of 0.1 in chloramphenicol supplemented LB-medium and 100 μL transferred into 96-well flat-bottom microplates. The wells were supplemented with AHL inducer (1:100 dilution) at appropriate concentrations and the cultures grown for an additional 3 hours in a shaking incubator at 37°C and 750rpm before measurement.

### Data analysis

Samples from cultured *E. coli* cells transformed with the composite devices plasmid constructs were analysed by flow cytometry (see Supplementary Information). Data analysis was carried out using GraphPad Prism^®^. For each of the AHL-receiver device, the mean cell fluorescence was calculated from 3 biological repeats. To determine GFP output, the mean cell autofluorescence value of *E. coli* TOP10 cells was subtracted from the mean cell fluorescence of AHL-receiver composite devices. The resulting values were fitted with a four-parameter logistical curve model (Equation 1). The fitting of the model was constrained with basal expression GFP output values derived from samples treated with 1*10^−15^ inducer concentration of the cognate AHL inducer. Fold activation for each device was calculated by dividing the maximal GFP fluorescence values by the basal GFP fluorescence values obtained from simulations of the fitted model where the devices showed GFP output saturation. Otherwise, fold activation was determined using the GFP output value at 100μM inducer concentration (highest soluble concentration under the experimental conditions). Activity of AHL-receiver devices was calculated by normalising their GFP output to the GFP output of the J23101 promoter. The same methodology was used to determine the relative activity of members of the Anderson library of constitutive promoters.

### Co-culture assay

Single colonies of *E. coli* Top10 cells transformed with two plasmid constructs (a quorum sensing composite device with a GFP reporter module, and a reporter plasmid that constitutively expressed another fluorescent protein as described in main text) were cultured overnight in LB-medium supplemented with chloramphenicol (34μg/ml) and ampicillin (50μg/ml), and cultured as previously described. A spectrophotometer was used to determine the OD_600_ value of cell cultures. This information was used to dilute cell cultures to an OD_600_ of 0.1 in antibiotic supplemented LB-medium, and these were then added together as a co-culture. A volume of 100 μl co-culture was transferred into 96-well flat-bottom microplate wells and induced with AHL at appropriate concentrations as defined in the main text. Finally, the microplate co-cultures were allowed to grow for 3 hours in a shaking incubator at 37°C and 750rpm. Co-culture samples were analysed using a BD LSRFORTESSA X-20 flow cytometer (BD Biosciences). Initially, cell populations were gated for blue fluorescence (excitation (ex): VL405nm/emission (em): VL450/50) and orange-red fluorescence (ex: YGL488nm/em: YGL586/15), and subsequently the orange-red population was gated for orange (ex: YGL561nm/em: YGL586/15) and red fluorescence (ex: YGL561nm/em: YGL670/30). The gated populations were subsequently analysed for green cell fluorescence (ex: BL488nm/em: YGL525/20). Analysis of mean cell fluorescence data for fold changes in GFP output was carried out as previously described. Analysis of flow cytometry data was carried out using the FlowJo™ software.

### Software tool

The software tool was developed in MATLAB from MathWorks^®^. It uses a database of the parameters from the fitted model input/output curves to calculate GFP output of AHL-receiver devices with all AHL chemical inducers combinations. The software then randomly selects a number of devices from the set of devices (equal to the number of communication channels specified by the user) and determines whether these devices satisfy the user specified conditions for activation and chemical crosstalk thresholds. In the case that the user specified condition are met, the program outputs the identified devices and the AHL inducer concentrations for which user-specified conditions hold true (0 = false, 1 = true). Alternative, if the randomly selected devices do not satisfy the conditions set by the user, the program randomly selects a different set of devices to test. This process is repeated until the user-defined conditions are met.

## AUTHORS CONTRIBUTIONS

Study design: NK, GBS and KP; Execution of experiments: NK; Data analysis: NK; Algorithm development: NK; Writing manuscript: NK; Editing manuscript: GBS, KP;

